# Optimization of mitochondrial membrane potential contributes to extended longevity of *daf-2* mutants in *C. elegans*

**DOI:** 10.64898/2026.11.18.767712

**Authors:** Aura A. Tamez González, Maisha Maliha Promi, Annika Traa, Jeremy M. Van Raamsdonk

## Abstract

Mitochondrial membrane potential is the charge difference across the inner mitochondrial membrane that is generated by the electron transport chain pumping protons from the mitochondrial matrix to the intermembrane space and is required for ATP synthesis, importing proteins into the mitochondria and movement of ions across the inner mitochondrial membrane. While a number of long-lived genetic mutants in *C. elegans* have been reported to have reduced mitochondrial membrane potential, increased mitochondrial membrane potential has also been associated with increased longevity. In this work, we examined the role of mitochondrial membrane potential in the long lifespan of *daf-2* insulin/IGF-1 receptor mutants. We found that mitochondrial membrane potential exhibits a small but significant decrease in day 1 *daf-2* mutants but a much greater decrease in day 8 *daf-2* worms. We also examined the effect of genes and compounds that have been shown to alter mitochondrial membrane potential on *daf-2* lifespan. Disruption of *ucp-4*, which increases mitochondrial membrane potential, decreased *daf-2* lifespan without reducing wild-type longevity. Treatment with the mitochondrial uncoupling compound FCCP (carbonyl cyanide-p-trimethoxyphenylhydrazone), which decreases mitochondrial membrane potential, also decreased the longevity of *daf-2* worms with no effect on wild-type lifespan. Combined, these results suggest that the altered mitochondrial membrane potential in *daf-2* mutants contributes to their extended longevity and that their long lifespan is dependent on maintaining mitochondrial membrane potential within an optimum range.

**Graphical abstract:** 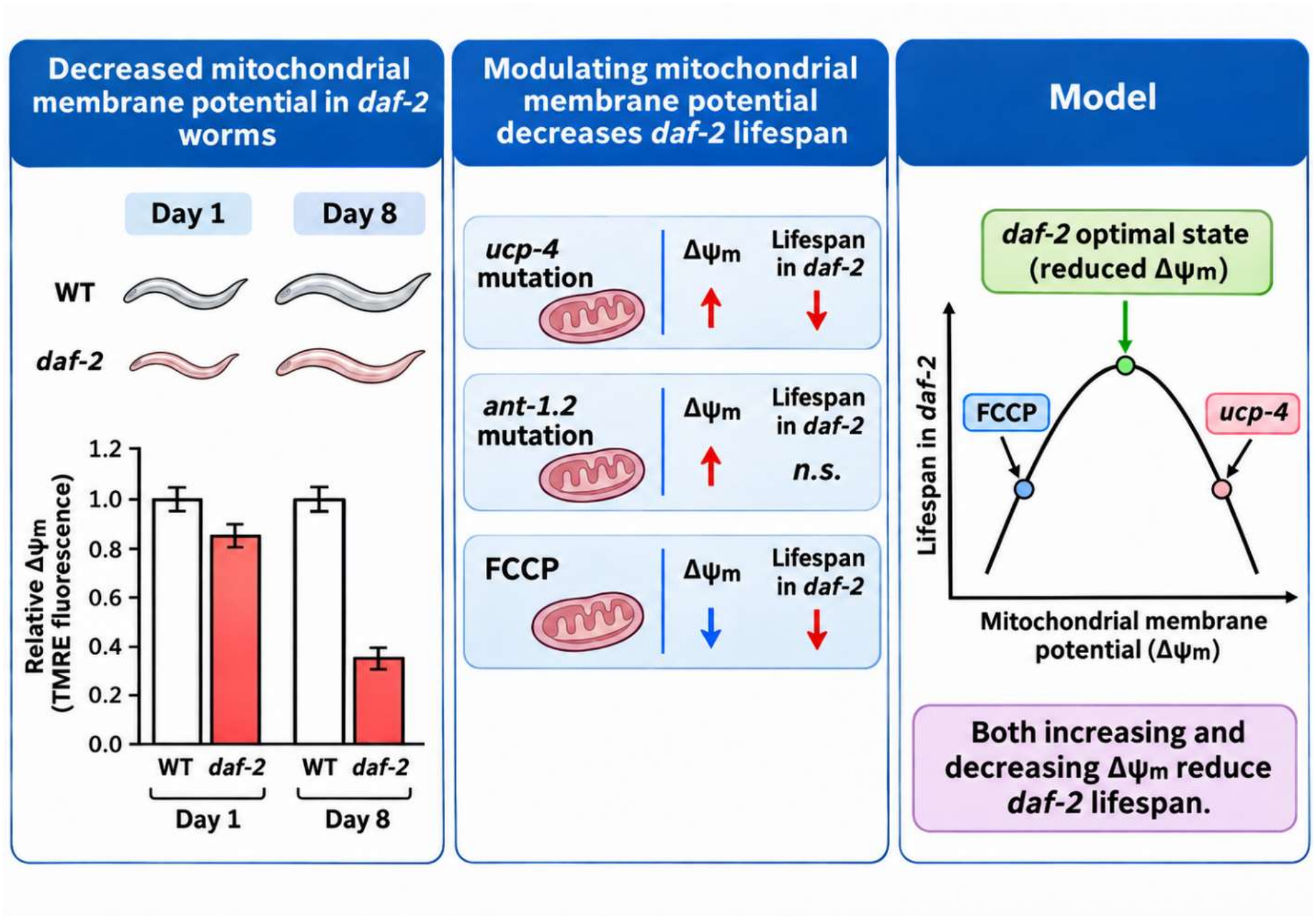

**Article Summary:** Mitochondria are a hub for multiple cellular functions and have an important role in lifespan determination. However, the effect of mitochondrial membrane potential on longevity is incompletely understood. We investigated the role of mitochondrial membrane potential in the long lifespan of *daf-2* insulin/IGF-1 receptor mutants in *C. elegans*. We found that mitochondrial membrane potential is decreased in *daf-2* mutants and that genes and compounds that either increase or decrease mitochondrial membrane potential specifically decrease *daf-2* lifespan. This suggests that reduced membrane potential in *daf-2* mutants contributes to longevity and that maintaining it within an optimal range is important for maximal lifespan.

## Introduction

Mitochondria perform a multitude of important functions within the cell. In addition to their role in producing energy in the form of ATP, mitochondria have important roles in signaling, metabolism, apoptosis and other cellular functions. In addition, mitochondria have been shown to affect longevity. While severe impairment of mitochondrial function shortens lifespan, a mild impairment of mitochondrial function has been shown to extend longevity. For example, mutations in genes that encode components of the mitochondrial electron transport chain, including *clk-1, isp-1* and *nuo-6* have been shown to increase lifespan (Lakowski and hekimi 1996; Feng *et al*. 2001; Yang and hekimi 2010). Similarly, knocking down the expression of a number of different genes expressing mitochondrial proteins results in increased lifespan (Dillin *et al*. 2002; Lee *et al*. 2003). In addition, recent work has demonstrated the importance of mitochondrial dynamics for longevity (Burkewitz *et al*. 2015; Weir *et al*. 2017; Traa *et al*. 2024a; Traa *et al*. 2024b).

Multiple previous studies have specifically examined the relationship between mitochondrial membrane potential and lifespan. Mitochondrial membrane potential is the electrochemical gradient across the inner mitochondrial membrane which is produced by the mitochondrial electron transport chain pumping protons from the mitochondrial matrix to the intermembrane space at mitochondrial complexes I, III and IV. This electrochemical gradient, known as the proton motive force, drives ATP synthesis via complex V ATP synthase. It was previously reported that a number of long-lived genetic mutants, including *daf-2, age-1, clk-1, isp-1* and *eat-2* worms all exhibit decreased mitochondrial membrane potential (Lemire *et al*. 2009). In that work, they also showed that RNAi knockdowns of 17 different genes that had previously been shown to increase lifespan also resulted in decreased mitochondrial membrane potential, while decreasing mitochondrial membrane potential in wild-type worms through treatment with the chemical uncoupler CCCP (carbonylcyanide-3-chlorophenylhydrazone) was sufficient to increase lifespan (Lemire *et al*. 2009).

While this work suggested that a decreased mitochondrial membrane potential is associated with or contributes to extended longevity, subsequent research suggests that the relationship is more complex. In wild-type worms, mitochondrial membrane potential declines with age (Brys *et al*. 2010; Gaffney *et al*. 2018; Berry *et al*. 2023a). Increasing mitochondrial membrane potential optogenetically during adulthood using a light-activated proton pump was shown to increase lifespan (Berry *et al*. 2023b) Similarly, dietary restriction or disruption of *eat-2* results in increased mitochondrial membrane potential and increased lifespan (Berry *et al*. 2023a). In addition, multiple other interventions, including treatment with Vitamin K, galactose, 2-deoxylucose or *Lactobacillus gasseri*, increase both lifespan and mitochondrial membrane potential (Nakagawa *et al*. 2016; Ali *et al*. 2025; Guo *et al*. 2026). At the same time, treatment with glucose or paraquat results in decreased mitochondrial membrane potential and decreased lifespan (Dilberger *et al*. 2019; Ali *et al*. 2025).

Combined, these results suggest that both increased or decreased mitochondrial membrane potential are compatible with extended longevity. In support of this conclusion, it was found that both short-lived (*gas-1, mev-1*) and long-lived (*isp-1, sod-2*) mitochondrial mutants have decreased mitochondrial membrane potential (Dingley *et al*. 2010). Finally, it was found that decreasing mitochondrial membrane potential through treatment with FCCP prevents the lifespan increase resulting from dietary restriction or disruption of *eat-2* but does not impact wild-type longevity (Berry *et al*. 2023a).

In this work, we further explored the relationship between mitochondrial membrane potential and lifespan, specifically focusing on *daf-2* insulin/IGF-1 receptor mutants. We find that mitochondrial membrane potential is decreased in *daf-2* mutants and that genes and compounds that modulate mitochondrial membrane potential decrease *daf-2* lifespan. This suggests that mitochondrial membrane potential in *daf-2* worms contributes to their lifespan extension and is optimized for their longevity.

## Results

As mitochondrial membrane potential has been associated with longevity, we quantified mitochondrial membrane potential in *daf-2* worms at day 1 and day 8 of adulthood. We found that mitochondrial membrane potential showed a small but significant decrease at day 1 of adulthood in *daf-2* mutants, while a much larger decrease was observed in day 8 *daf-2* worms (**Figure 1**).

**Figure 1.**
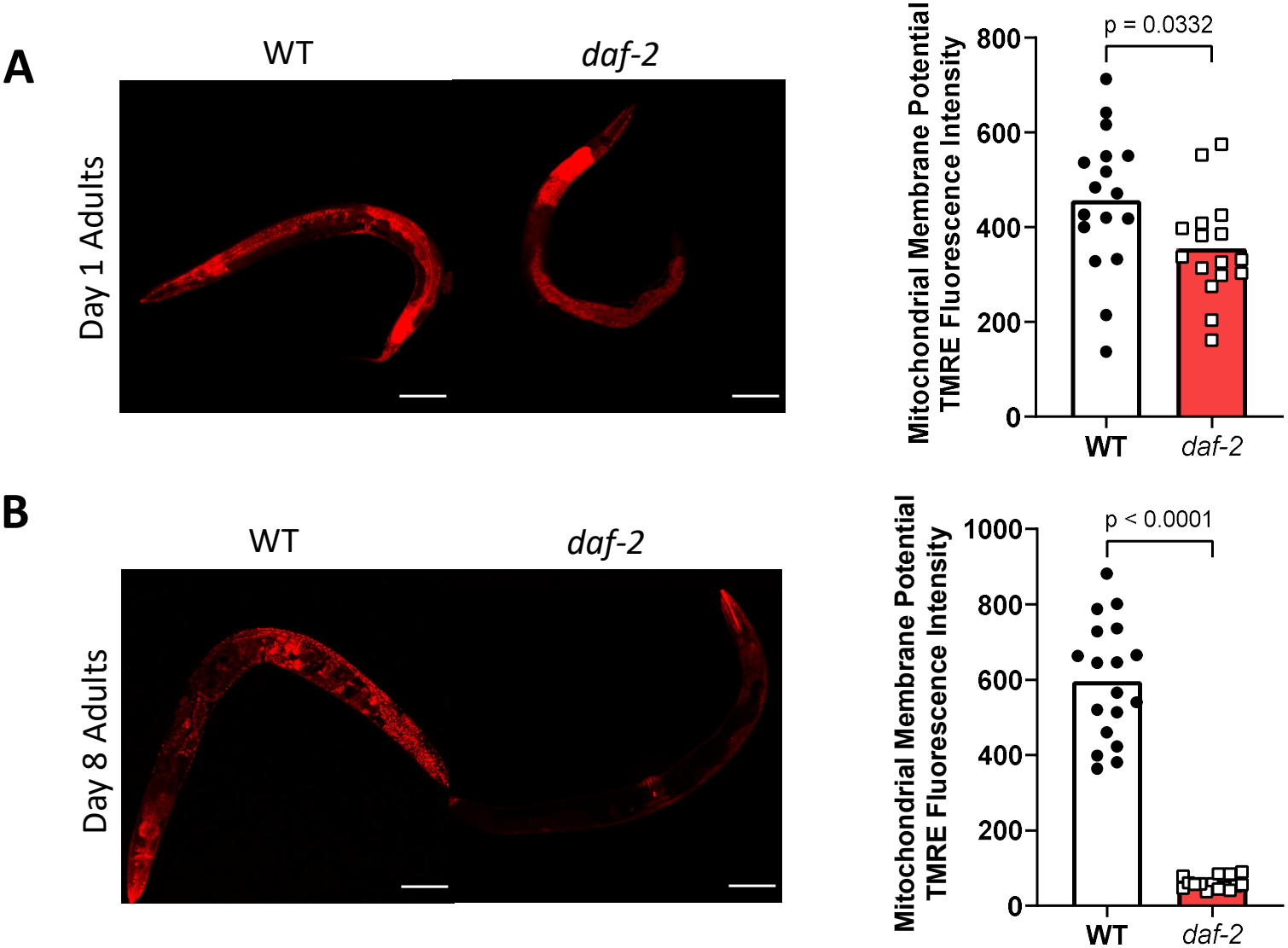
Long-lived *daf-2* insulin/IGF-1 receptor mutants have decreased mitochondrial membrane potential. Worms at day 1 and day 8 of adulthood were stained with tetramethylrhodamine ethyl ester (TMRE) to measure mitochondrial membrane potential. (**A**) At day 1 of adulthood, *daf-2* worms exhibit a small but significant decrease in mitochondrial membrane potential compared to wild-type worms. (**B**) By day 8 of adulthood, *daf-2* worms have a markedly decreased mitochondrial membrane potential compared to wild-type worms. Statistical significance was assessed using a t-test. Scale bar indicates 100 µm.

Based on the observation that mitochondrial membrane potential is altered in *daf-2* worms, we sought to determine the extent to which this altered membrane potential contributes to their extend longevity. Accordingly, we treated *daf-2* worms with genetic mutations or compounds that have been shown to modulate mitochondrial membrane potential and measured lifespan. First, we crossed *daf-2* mutants to *ucp-4* and *ant-1*.*2* mutants, which have previously been shown to increase mitochondrial membrane potential (Berry *et al*. 2023a). We found that disruption of *ucp-4* significantly decreased *daf-2* lifespan with no detrimental effect on wild-type longevity (**Figure 2A**). Deletion of *ant-1*.*2* did not significantly affect the lifespan of wild-type of *daf-2* worms (**Figure 2B**).

**Figure 2.**
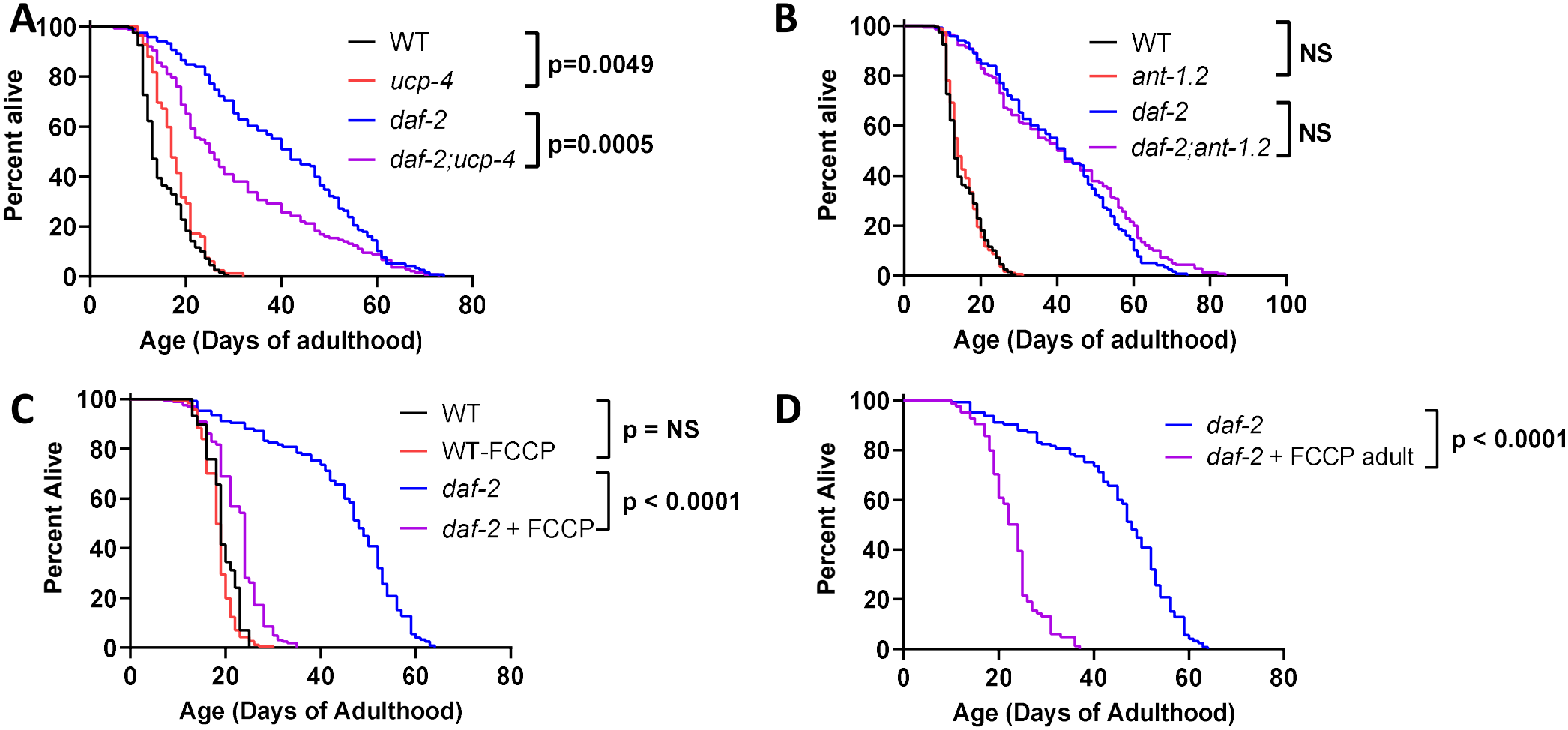
Genes and compounds that alter mitochondrial membrane potential decrease *daf-2* lifespan. To examine the effect of altering mitochondrial membrane potential on *daf-2* lifespan, *daf-2* worms were either crossed to *ucp-4* and *ant-1*.*2* mutants or treated with the uncoupling compound carbonyl cyanide-p-trimethoxyphenylhydrazone (FCCP) (**A**) Disruption of *ucp-4* decreased *daf-2* lifespan despite mildly increasing lifespan in wild-type worms. (**B**) Deletion of *ant-1*.*2* did not affect lifespan in wild-type or *daf-2* worms. (**C**) Life-long treatment with FCCP markedly decreased the lifespan of *daf-2* worms but did not affect wild-type lifespan. (**D**) Treatment of *daf-2* worms with FCCP beginning at adulthood also markedly decreases their longevity indicating that the effect of FCCP on *daf-2* lifespan is not developmental. Three biological replicates were performed. Statistical significance was assessed using a log-rank test.

We next treated *daf-2* worms with the uncoupling compound carbonyl cyanide-p-trimethoxyphenylhydrazone (FCCP) to decrease the mitochondrial membrane potential. FCCP has previously been shown to decrease mitochondrial membrane potential in *C. elegans* (Kwon *et al*. 2018; Berry *et al*. 2023a). We found that treating *daf-2* worms with 10 µM FCCP throughout their entire life markedly decreased their lifespan almost back to wild-type (**Figure 2C**). In contrast, FCCP treatment did not affect wild-type lifespan indicating that the chemical uncoupler specifically affects *daf-2* lifespan.

To avoid possible detrimental effects of manipulating mitochondrial membrane potential during development, we examined the effect of treating *daf-2* worms with FCCP during adulthood only. Similar to life-long treatment, we found that adult-only treatment with FCCP markedly decreased *daf-2* lifespan (**Figure 2D**). Together, these results suggest that the decreased mitochondrial membrane potential in *daf-2* worms is optimized for their longevity, as genes and interventions that shift mitochondrial membrane potential higher or lower both decrease *daf-2* lifespan.

## Discussion

Consistent with previous work, we show that mitochondrial membrane potential is decreased during adulthood in *daf-2* worms (Lemire *et al*. 2009). Expanding upon the earlier work, our results suggest that the magnitude of decrease in mitochondrial membrane potential is different at different ages with a larger decrease being observed at day 8 compared to day 1 of adulthood. Importantly, our work demonstrates the importance of the lower mitochondrial membrane potential for the long lifespan of *daf-2* worms as genes and interventions that further modulate mitochondrial membrane potential decrease *daf-2* lifespan suggesting that the level of decrease is optimized for their longevity.

Collectively, our results combined with previous reports in the literature support a model in which mitochondrial membrane potential has a non-linear relationship with longevity. Both increased and decreased mitochondrial membrane potential are compatible with extended longevity, and experimentally increasing or decreasing mitochondrial membrane potential can both increase or decrease lifespan. Overall, it appears that there is an optimal mitochondrial membrane potential with respect to longevity and that it is dependent on genotype, age, and multiple other factors.

## Materials and Methods

### Strains

|  |  |
|---|---|
| N2 | WT |
| CB1370 | *daf-2(e1370) III* |
| VC620 | *ant-1.2(gk294) I* |
| MQ1625 | *ucp-4(ok195) V* |
| JVR716 | *daf-2(e1370) III; ant-1.2(gk294) I* |
| JVR717 | *daf-2(e1370) III; ucp-4(ok195) V* |

### Generation of double mutants

Double mutant strains (*daf-2(e1370);ant-1*.*2(gk294)* and *daf-2(e1370);ucp-4(ok195)*) were generated using standard genetic crossing techniques. First, N2 males were crossed with *daf-2* mutant hermaphrodites. Male progeny from this cross were subsequently crossed with either *ant-1*.*2* or *ucp-4* mutant hermaphrodites. Male progeny from the second cross were then backcrossed to the corresponding *ant-1*.*2* or *ucp-4* deletion mutants. The resulting hermaphrodite progenies were allowed to self-fertilize after which they were transferred to 25°C and allowed to lay eggs. After egg laying, adult worms were subsequently removed, and the eggs were maintained at 25°C to induce dauer formation in order to select for worms homozygous for the *daf-2* mutation. After 2–3 days, dauers were transferred to a new plate. After an additional day at 25 °C, worms that were still dauer were recovered at 16 °C, singled onto individual plates and their genotypes were determined by PCR using primers specific to the *ant-1*.*2* and *ucp-4* deletion alleles, followed by DNA sequencing to confirm the *daf-2* mutation. Animals homozygous for both mutations were selected to establish stable double-mutant strains for subsequent experiments.

### Mitochondrial membrane potential - Tetramethylrhodamine (TMRE) staining

The potentiometric fluorescent indicator, TMRE, was used to measure mitochondrial membrane potential. TMRE was dissolved in DMSO to make a 50 mM stock solution. The TMRE stock solution was then diluted in M9 to make a 5 μM working solution. The working TMRE solution was then pipetted onto NGM plates seeded with OP50 and were left on a nutator for 30 min to allow the TMRE to spread evenly over the agar. Approximately 20 age-matched day 1 or day 8 worms were transferred to the TMRE plates, which were then covered to protect from light and stored at 20 °C for 20 h. To de-stain, worms were transferred to regular NGM plates seeded with OP50 and left at 20°C for 4 h. Worms were imaged with the 20× objective using a Zeiss LSM 780 confocal microscope and ImageJ was used to quantify the fluorescence intensity of TMRE in the whole body of the worm.

### Lifespan

Lifespan assays were conducted at 20°C on plates containing 50 μM 5-fluoro-2′-deoxyuridine (FUdR) to limit the growth of progeny. This concentration was previously shown to have minimal effects on the longevity of wild-type worms (VAN RAAMSDONK AND HEKIMI 2011). For the FCCP experiment, FCCP was added to the NGM media prior to pouring plates,to a final concentration of 10 µM. Lifespan plates were seeded with OP50 bacteria and were kept at 20°C. Viability was assessed every 2-3 days by gentle prodding with a platinum pick. Worms that either had internal hatching of progeny or expulsion of internal organs were censored and not counted as deaths. At least three biological replicates with 30-40 worms per replicate were completed for each strain and condition.

### Statistical analysis

Statistical significance of differences between groups was determined by computing a t-test or a log-rank test using Graphpad Prism, as indicated in the figure legends. All error bars indicate the standard error of the mean.

## Acknowledgments

Some strains were provided by the CGC, which is funded by NIH Office of Research Infrastructure Programs (P30 OD010440). We would also like to acknowledge the *C. elegans* knockout consortium and the National Bioresource Project of Japan for providing strains used in this research. This work was supported by the Canadian Institutes of Health Research (CIHR; http://www.cihr-irsc.gc.ca/ ; JVR) and the Natural Sciences and Engineering Research Council of Canada (NSERC; https://www.nserc-crsng.gc.ca/index_eng.asp; JVR). The funders had no role in study design, data collection and analysis, decision to publish, or preparation of the manuscript. ChatGPT version GPT-6 was used to generate the graphical abstract. The graphical abstract was checked and revised until it was scientifically correct and appeared as envisioned and described.

## Author Contributions

Conceptualization: JVR. Methodology: ATG, MMP, AT, JVR. Investigation: ATG, MMP, AT, JVR. Analysis: ATG, MMP, AT, JVR. Validation: ATG, MMP, AT, JVR. Visualization: ATG, MMP, AT, JVR. Writing – original draft: JVR. Writing – review and editing: ATG, MMP, AT, JVR. Supervision: JVR.

## Data and materials availability

Raw data lifespan data is provided in Table S1. All materials used in this manuscript are available to be shared with the scientific community. Requests for data or materials should be addressed to Jeremy Van Raamsdonk.

